# Ghost lineages highly influence the interpretation of introgression tests

**DOI:** 10.1101/2021.03.30.437672

**Authors:** Théo Tricou, Eric Tannier, Damien M. de Vienne

## Abstract

Most species are extinct; those that are not are often unknown. Sequenced and sampled species are often a minority of known ones. Past evolutionary events involving horizontal gene flow, such as horizontal gene transfer, hybridization, introgression and admixture, are therefore likely to involve “ghosts”, *i.e*. extinct, unknown or unsampled lineages. The existence of these ghost lineages is widely acknowledged, but their possible impact on the detection of gene flow and on the identification of the species involved is largely overlooked. It is generally considered as a possible source of error that, with reasonable approximation, can be ignored. We explore the possible influence of absent species on an evolutionary study by quantifying the effect of ghost lineages on introgression as detected by the popular D-statistic method. We show from simulated data that under certain frequently encountered conditions, the donors and recipients of horizontal gene flow can be wrongly identified if ghost lineages are not taken into account. In particular, having a distant outgroup, which is usually recommended, leads to an increase in the error probability and to false interpretations in most cases. We conclude that introgression from ghost lineages should be systematically considered as an alternative possible, even probable, scenario.

## 1. Introduction

Evolutionary studies are always restricted to a subset of species, populations or individuals. This is by choice, because only a fraction of the data is relevant to the question being addressed, and by necessity, because the approaches used have methodological and technical limitations. Another reason is that most lineages are simply unknown. More than 99.9% of all species that have ever lived are now extinct (Raup 1991) and only a small fraction of extant species have been described. The number of extant eukaryote species that are still uncatalogued is almost an order of magnitude higher than the number of those reported (~1.3 million species have been catalogued, Mora et al. 2011), and is many orders of magnitudes higher if we consider Bacteria and Archaea diversity (Locey and Lennon 2016).

Taking these extinct, unknown or unsampled “ghost” lineages into account is particularly important when studying introgression, *i.e*. the integration of genetic material from one lineage to another *via* hybridization and subsequent backcrossing. This mode of gene flow across species boundaries appears to be common in the Eukaryotic domain and has been shown to be adaptive in some cases (see for example Hedrick 2013 for a review). Introgression has been reported in such diverse lineages as humans (Green et al. 2010; Meyer et al. 2012), boars (Liu et al. 2019), butterflies (Martin et al. 2013; Smith and Kronforst 2013; Massardo et al. 2020), fishes (Schumer et al. 2016; Meier et al. 2017), plants (Eaton and Ree 2013; Zhang et al. 2019) and fungi (Zhang et al. 2018; Keuler et al. 2020), to name but a few. Since ghost lineages are probably massively present around any phylogeny of extant species, many gene flow events that are detectable now are likely to have involved a ghost lineage. This has been repeatedly acknowledged (Maddison 1997; Galtier and Daubin 2008; Green et al. 2010; Eaton and Ree 2013; Szöllősi et al. 2013, 2015), especially in studies of introgression between populations, but it was considered either a source of noise (Pease and Hahn 2015), or a problem that could be resolved by adding new species as they become available, or by combining the results of multiple detection tests (Eaton et al. 2015; Kumar et al. 2017; Barlow et al. 2018). Recently, Hibbins and Hahn (2021) advised bearing ghost lineages “in mind” when investigating gene flow but, as far as we know, the real impact of ghost lineages on the ability of different methods to detect gene flow and correctly identify involved lineages has not been properly evaluated and quantified.

Over the past few years, the ever-growing number of sequenced genomes and the development of new methods have improved the detection of introgression. One of the most widely used methods for inferring introgression is the D-statistic (or Patterson’s D), also known as the ABBA-BABA test (Kulathinal et al. 2009; Green et al. 2010; Durand et al. 2011; Patterson et al. 2012). There are many reasons for its success. The D-statistic is easy to understand and implement, quick to compute and easy to interpret. This method is based on phylogenetic discordance and can discriminate incongruence caused by Incomplete Lineage Sorting (ILS) from incongruence caused by gene flow (Kulathinal et al. 2009; Green et al. 2010; Durand et al. 2011; Patterson et al. 2012). The ABBA-BABA test considers four taxa: three ingroup taxa and one outgroup, with a ladder-like phylogenetic relationship (Fig. 1). The test relies on counts of the number of sites that support a discordant topology. Two biallelic SNP patterns are considered, ABBA and BABA, depending on which allele (A: ancestral, B: derived) is present in each taxon. The D-statistic is computed using the classic formula from Durand et al. (2011):

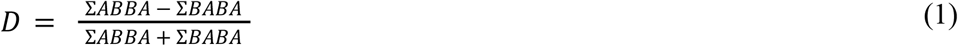

The null hypothesis states that under a scenario with no gene flow, both ABBA and BABA patterns can be attributed to ILS and thus should be observed in equal numbers. Significant deviation from this expectation, resulting in a D-statistic significantly different from zero, is usually interpreted as introgression between two of the three lineages forming the ingroup (Fig. 1, left panel). The outgroup should be distant enough from the ingroup such that it is not involved in an introgression with any of the ingroup lineages (Green et al. 2010; Osborne et al. 2016; Irwin et al. 2018).

**Figure 1.**
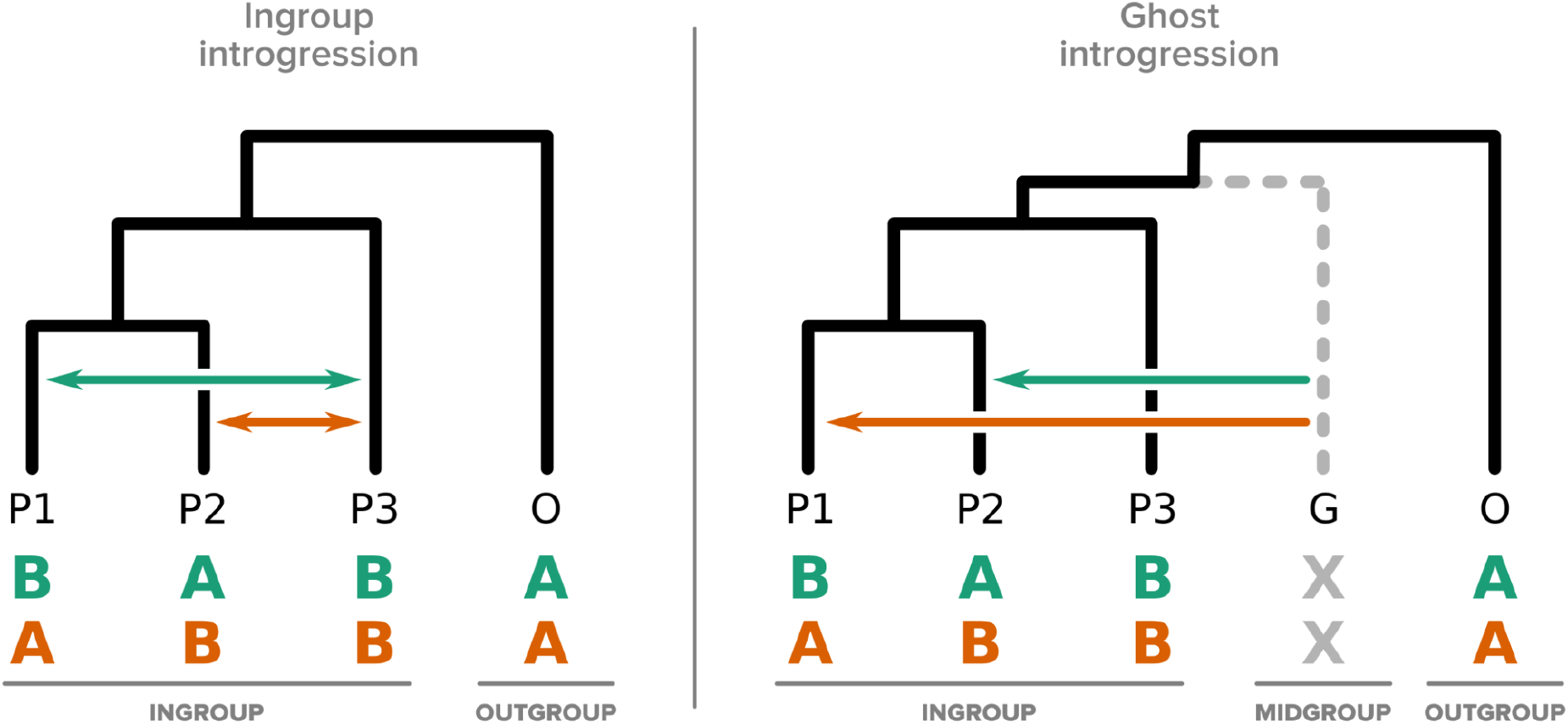
Introgression events that can result in a significant excess of ABBA or BABA patterns according to the D-statistic. The usual interpretation of this excess is the hypothesis of “ingroup” introgression (left panel). However “ghost” (or “midgroup”) introgression (right panel) from ghost lineages (G) can produce similar patterns.

Undersampling is known to be one of the factors that can possibly confound the D-statistic (Martin et al. 2015; Zheng and Janke 2018), and affect the detection of introgression. This is because using a subset logically leads to an underestimation of the true frequency of introgression and thus inflates the role of ILS (Maddison and Knowles 2006). It has been clearly stated that the donor genome could easily be misidentified because introgression from a sampled lineage *(e.g*. P3) or from one sister ghost lineage to the same recipient lineage would produce the same signal and result in indistinguishable D-statistic results (Eaton and Ree 2013; Eaton et al. 2015; Pease and Hahn 2015; Zheng and Janke 2018). Another stronger impact of ghost lineages, however, was foreseen early on in the history of the test (by Durand et al. (2011), in their first description of the test), but has been largely overlooked afterwards: introgression from a ghost lineage between the ingroup and the outgroup (the “midgroup”, see Fig. 1) could lead to the wrong identification of both the donor ***and*** the recipient genomes (Fig. 1, right panel). Under this scenario, none of the species thought to be involved in the introgression event are correctly identified. This possible source of error in the interpretation of the D-statistic has often been acknowledged (Durand et al. 2011; Ottenburghs et al. 2017;

Zheng and Janke 2018; Hibbins and Hahn 2021) but surprisingly, it does not seem to have changed the way the test is commonly interpreted, perhaps because it is thought that the impact of this possibility is low, even though it has not been formally quantified. This is the goal of this study.

We begin by an illustration of the possibility of misinterpreting the ABBA-BABA test when some species are unknown or not included using a previously published bear phylogeny, recurrently used later on to estimate parameters on realistic situations. We then quantify the effect of ghost lineages on the misidentification of the donor and the recipient lineage using simulations. We explore the impact of outgroup choice, number of unsampled species and genetic divergence between introgressed taxa on the probability of misinterpreting introgression events.

We show that under the realistic assumption that there are many ghost lineages branches in the tree, and assuming a simple demographic history of the populations considered, most significant D-statistics are attributable to ghost lineages. This suggests that most of the lineages involved (donors and recipients) are incorrectly identified by the usual interpretation of D-statistics. The error rate increases with the distance between ingroup and outgroup, even though the outgroup is usually chosen so that its distance from the ingroup is sufficient to avoid any introgression between the two (Green et al. 2010; Osborne et al. 2016; Irwin et al. 2018). This observation, that a close outgroup as well as a distant outgroup is a source of interpretation error, hampers the delimitation of a safe zone for the interpretation of the D-statistic.

These results call for a new way of interpreting D-statistics, and more generally call into question established methods of introgression detection. Our results illustrate the recent statement by Ottenburghs (2020) that “the presence of ghost introgression has important consequences for the study of evolutionary processes”, and provide a demonstration of this importance.

## 2 Materials and Methods

### 2.1 Bear genomic dataset

We use the dataset from Barlow et al. (2018) to illustrate the possibility of misinterpreting significant results of an ABBA-BABA test. This dataset has the advantage of being easily available and to support, according to the authors, a simple introgression scenario, *i.e*. a documented introgression between polar bears and brown bears from the ABC Islands in Alaska. We downloaded the genome sequences (Barlow et al. 2019) of three brown bears (*Ursus arctos*) from Alaska (id: Adm1), Russia (id: 235) and Slovenia (id: 191Y), one polar bear (*U. maritimus;* id: NB) and one American black bear (*U. americanus;* id: Uamericanus), all aligned against the panda reference genome (Li et al. 2010). Their relationship is ((((Alaska = P1, Russia = P2), Slovenia = P3), Polar bear = P4), Black bear = O). Using scripts available from the GitHub repository of Barlow et al. (2018) https://github.com/jacahill/Admixture, we computed the D-statistic for two quartets: (((Alaska,Russia),Polar bear),Black bear) and (((Alaska,Russia),Slovenia),Black bear). We used the script for all sites (not transversions only like in Barlow et al, 2018) as we do not have any archaic species in the dataset. The weighted block jackknife from the script was used to compute the Z-score in non-overlapping 1 Mb windows with a script available on the GitHub repository mentioned above. We considered the result to be significant if it was more than three standard deviations from zero (Z > 3 or Z < −3), as *per* Green et al. (2010).

### 2.2 Species tree and gene tree simulation

Species trees were simulated using the birth-death simulator implemented in the R function *rphylo* from the *ape* package (Paradis and Schliep 2019). Speciation rate was fixed at 1, extinction rate at 0.9 and the simulation was stopped when N extant lineages, varying in {20, 40, 60, 80, 100}, were present in the species tree (step 1 in Figure 2). Then 20 taxa were uniformly sampled from the N taxa. An introgression event was chosen in the species tree (including unsampled lineages), with a donor and a recipient. The donor branch was selected among the branches of the species tree with a probability proportional to its length, and the time of introgression uniformly at random on this branch. The recipient was randomly sampled among the lineages present at the time of the introgression, with a probability that decreases exponentially with the phylogenetic distance from the donor (step 2 in Figure 2):

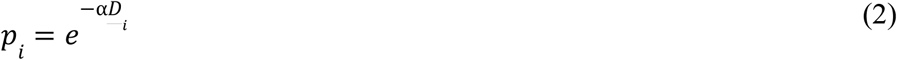

**Figure 2.**
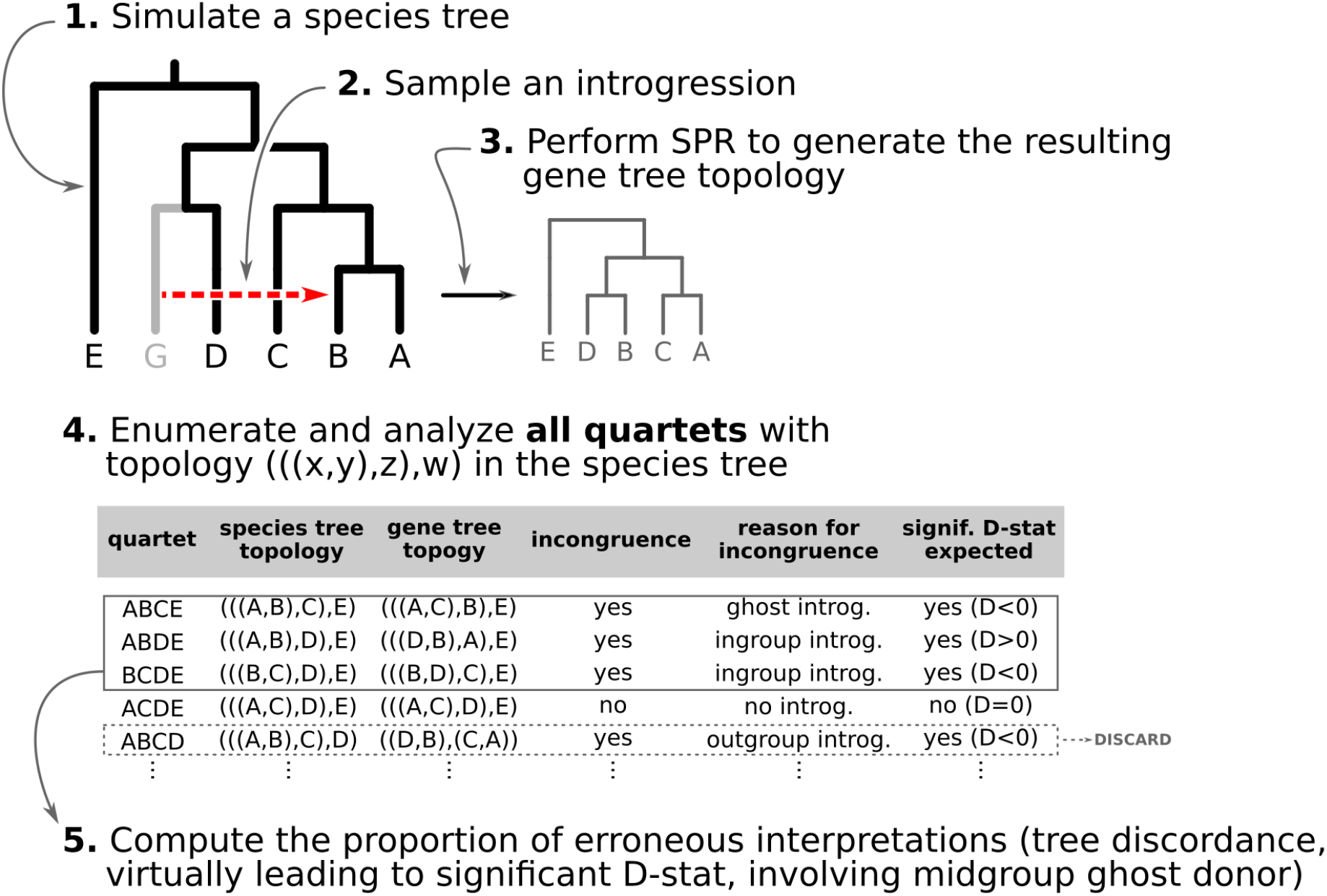
Species tree/Gene tree simulation: (1) a species tree is generated under a birth death model and 20 taxa are sampled from it; (2) an introgression event is picked from a random donor and recipient; (3) an introgressed gene tree is constructed from the species tree by SPR; (4) for each quartet with a ladder-like topology (((x,y),z),w) in the species tree, species tree and gene tree topologies are compared to determine if there is an incongruence caused by the introgression; (5) the proportion of erroneous interpretation of the D-statistic across the species tree is computed by the sum of all introgressions with a midgroup ghost donor over all introgressions detected, outgroup introgressions excluded.

Where 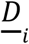 is the distance from the *i*-th recipient, normalized by the distances from all possible recipients, and α is a parameter in {0, 1, 10, 100, 1000}. With *α* = 0, introgressions occur between any contemporaneous branches on a tree with equal probability. When *α* increases, introgressions are more likely to occur between closely related taxa. If the recipient is a branch with no extant offspring, then the introgression cannot be detected, so only introgressions such that the recipient has descendants among the 20 remaining species were kept. After setting the donor and recipient, a gene tree was generated with subtree pruning and regrafting (SPR) (Bordewich and Semple 2005), simulating the introgression event (step 3 in Figure 2). All unsampled lineages are then pruned from species and gene trees.

### 2.3 Comparison of the species tree and the gene tree as proxy for the D-statistic

For each gene tree/species tree pair, we counted all species quartets with a ladder-like topology (((P1,P2),P3),P4) in the species tree and (((P1,P3),P2),P4) or (((P2,P3),P1),P4) in the gene tree. These configurations were interpreted as yielding significant D-statistics. This avoids the computational burden of simulating sequences for a high number (250 000, see Section 2.5) of cases and gives reasonably equivalent results (Supplementary Material Section 1).

We then counted the number of situations where the result was correctly interpreted (the simulated introgression is between extant lineages in this quartet) or misinterpreted (the simulated introgression is from ghost lineage) (step 5 in Figure 2).

### 2.4 Measuring the distance to the outgroup

For each species quartet we computed the distance between outgroup and ingroup using the ratio *R*=*t1*/*t2,* where *t1* is the distance (sum of branch lengths) between the most recent common ancestor of the ingroup and the most recent common ancestor of all four taxa (see *t1* in Fig. 5), and *t2* is the total height of the four-taxon tree (see *t2* in Fig. 5). To correlate this distance with the rate of interpretation error of D-statistics, we identified ten intersecting subsets of quartets according to their *R* value: subsets for which *R* is higher than a threshold “x”, with “x” varying from 0 to 0.9 with a step of 0.1. We computed the rate of interpretation error for each of the 10 subsets.

### 2.5 Simulation dataset

We simulated, for each value of N in {20, 40, 60, 80, 100} and *α* in {0, 1, 10, 100, 1000}, 100 species trees with N species, and for each species tree, 100 independent gene trees with independent introgression events. For each gene tree/species tree pair, 20 species were uniformly sampled from N (extant species), and the rest were pruned, resulting in 250,000 pairs of trees each with 20 leaves.

## 3 Results

### 3.1 Bear phylogeny exemplifies the problem of interpreting the D-statistic without taking unsampled lineages into account

Using genomic data, we show how the presence or absence of one lineage, in this case the polar bear, can lead to opposite interpretations of the D-statistic if interpreted without considering ghost lineages. From the bear phylogeny (Material and Methods; phylogeny shown in Figure 3A), we removed either the Slovenian bear (Figure 3B, subset 1), which is not thought to have introgressed with other bear species from the ingroup, or the polar bear (subset 2), which is suspected to have introgressed with brown bears from Alaska (Cahill et al. 2013; Liu et al. 2014; Lan et al. 2016; Kumar et al. 2017; Barlow et al. 2018). In the first subset, we identified 175,413 ABBA patterns and 226,992 BABA patterns resulting in a significant negative D-score (D = −0.128, Z = −11.71), which is congruent with introgression between polar bears and Alaskan bears (Fig. 3B) in the usual interpretation of the test. In the second subset, we identified 266,173 ABBA patterns and 213,830 BABA patterns resulting in a significant positive D-score (D = 0.109, Z = 14.24).

**Figure 3.**
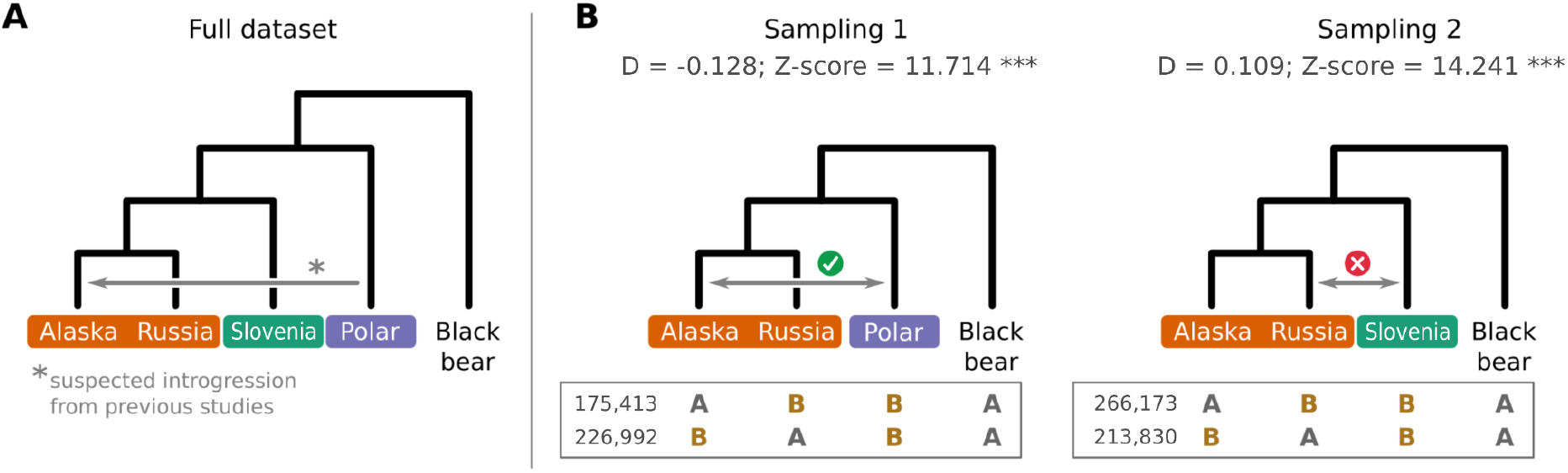
The effect of sampling on the interpretation of the D-statistic, using bear genomic data as an example. **A**. Phylogenetic relationship of the five bear taxa sampled. The grey arrow shows the introgression inferred from previous studies. **B**. D-statistic calculated from two four-taxon subsets. The number of ABBA and BABA patterns is given below the trees. In subset 1, the Slovenian bear, a lineage that is not thought to be involved in introgression, is removed. In subset 2, the donor of the introgression shown in 3A (*i.e*. the polar bear) is removed. Introgressions were inferred from the D-statistic (grey arrows), and their congruence with other studies (green tick = congruent, red cross = not congruent) is indicated above the arrow.

If we were to interpret this second result as evidence of introgression between the lineages sampled here (Alaskan, Russian, Slovenian and black bears), we would conclude that there was introgression between bears from Slovenia and Russia (Figure 3B), even though this significant positive D-statistic could also be attributed to introgression between polar bears (not sampled here) and Alaskan bears. The latter attribution, however, relies on our knowledge of the existence of polar bears, considered a ghost lineage. This hypothesis could similarly be called into question if we knew the existence of another lineage, because we can never assume that we know all the lineages leading to extant or extinct species. Thus, even with good taxonomic sampling, there is a real chance that an interpretation based only on known lineages wrongly infers introgression events.

### 3.2 Significant D-statistics are often due to introgressions from ghost lineages

Using simulated datasets (Material and Methods), we estimated the frequency of misinterpreting introgression events. We counted the number of D-statistics due to midgroup ghost introgressions (corresponding to the *proportion of erroneous interpretations)*. We observed between 15% and 100% of erroneous interpretations, the frequency of which increased with (i) the proportion of unsampled lineages, (ii) the distance between ingroup (P1, P2, P3) and outgroup O, and (iii) the probability of introgression between distantly related lineages (Supplementary Material Section 2 for a complete summary). We describe these three trends in detail in the following sections and relate the range of each parameter we used to biological data such as the bear genomes described above.

#### 3.2.1. The proportion of unsampled species

The effect of absent lineages on the interpretation of the D-statistic was investigated using simulated species trees with N = {20, 40, 60, 80, 100} from which 20 species were randomly sampled. This corresponds to a sampling effort ranging from 100% (20 species out of 20) to 20% (20 species out of 100). We observed that low sampling contributes to an increase in the number of misinterpreted D-statistics due to ghost introgression (Fig. 4). While the mean proportion of erroneous interpretations is ~25% when 100% of extant lineages are sampled, it is close to 60% when only 20% of extant lineages are sampled.

**Figure 4.**
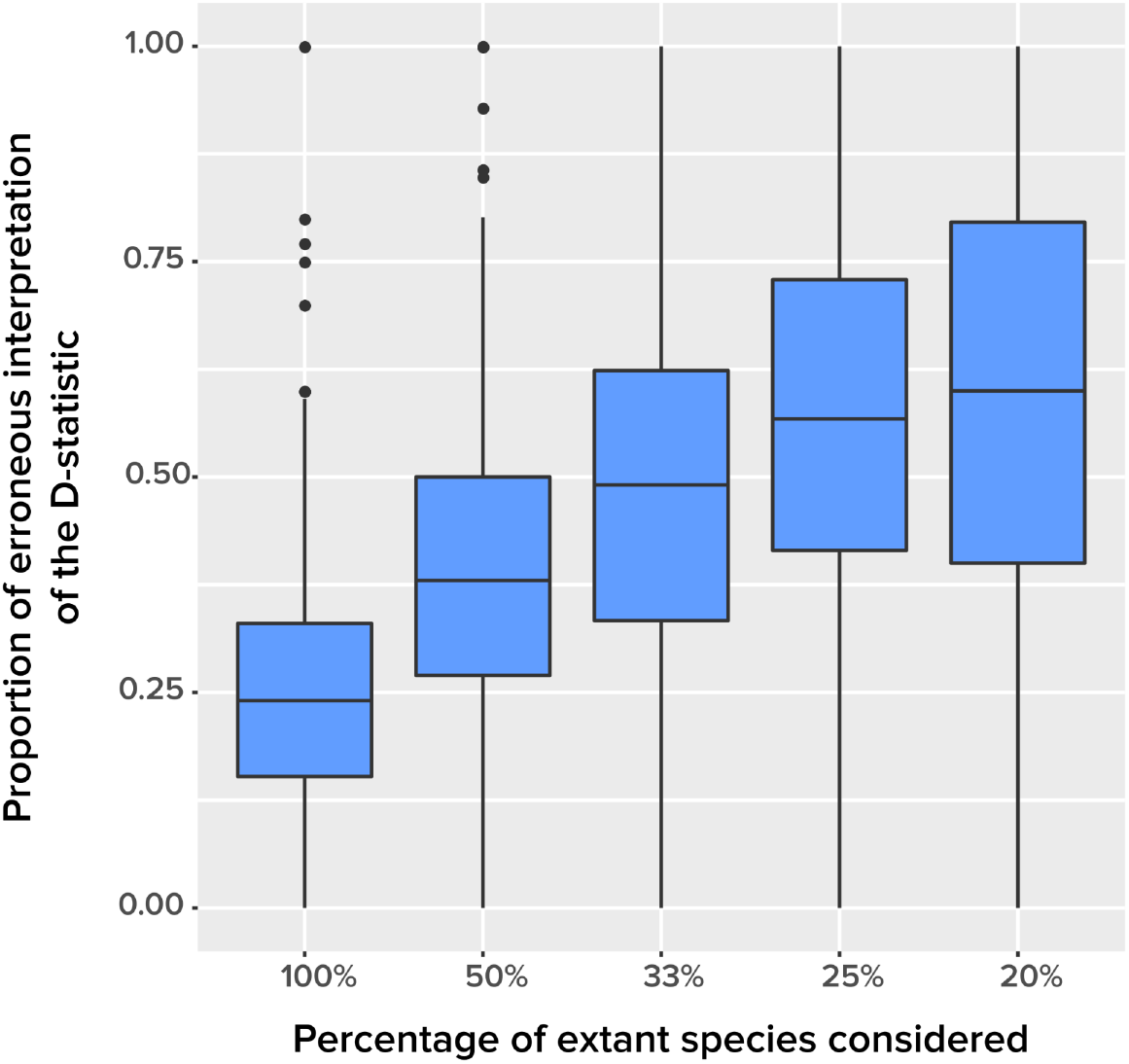
The effect of taxonomic sampling (x-axis) on the proportion of erroneous interpretations of the D-statistic (y-axis). The error rate is increasing with the amount of unknown.

For example, to study introgression in bears, Barlow et al. (2018) sampled 13 *Ursus* species and one *Ailuropoda* species, both members of the family Ursidae, based on the availability of genomic data. However, the Global Biodiversity Information Facility (gbif.org) reports 140 species in Ursidae, 100 of them belonging to the genus *Ursus*, which is close to the highest error rate in the simulations.

By contrast, and unlike we initially expected, we found no correlation between the number of extinct lineages in the tree and the proportion of erroneous interpretations of the D-statistic (Supplementary Material Section 3). Our interpretation is that increasing the number of extinct lineages is achieved by increasing the probability of extinction in the birth-death process, which also increases the probability that mid-group lineages, the possible source of ghost introgressions, become extinct before having the opportunity to introgress. Further investigations are needed to better characterize this effect.

#### 3.2.2 The distance between outgroup and ingroup

In ABBA-BABA tests, the outgroup is usually chosen so that its distance from the ingroup is sufficient to minimize the chance of introgression between the two (Green et al. 2010; Osborne et al. 2016; Irwin et al. 2018). Zheng and Janke (2018) stated that the distance between outgroup and ingroup had little to no impact on the sensitivity of the D-statistic. However, they focused on evaluating the effect of saturation of sequence substitutions in the outgroup and did not consider possible introgressions from mid- or outgroups.

From our simulations, we observed that the proportion of ghost introgressions (leading to erroneous interpretations) increased with *R*, the relative distance to the outgroup (see Materials and Methods). On average, when *R* > 0.3, more than 50% of the significant D-statistics are associated with ghost introgressions (Fig. 5). We found that, when *R* > 0.7, a median of 100% of D-statistics resulted from ghost introgressions.

**Figure 5.**
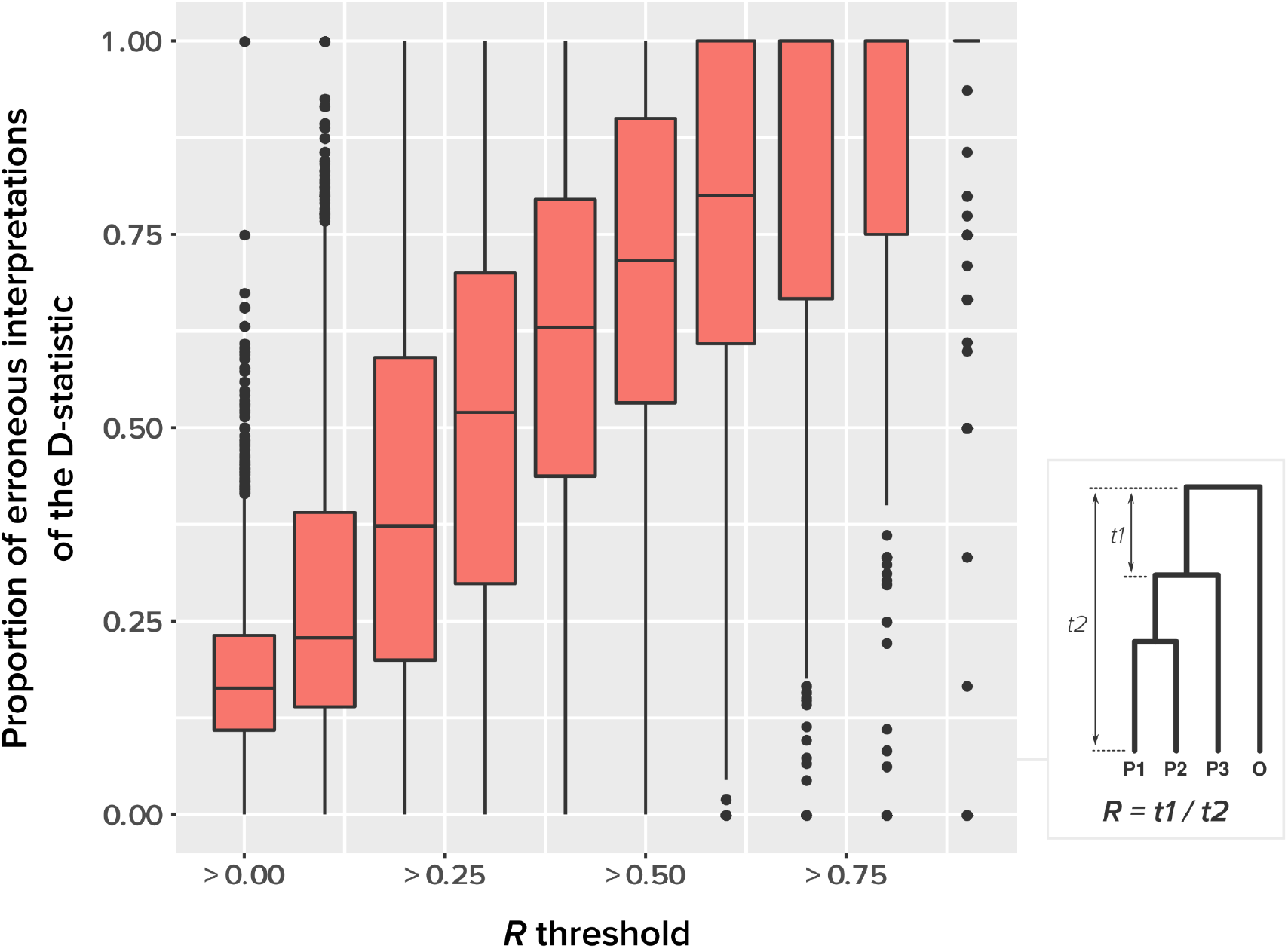
The relationship between outgroup distance (*R*) and the proportion of erroneous interpretations of the D-statistic for different thresholds of *R* (x-axis). The distances *t1* and *t2* used to calculate the relative distance to the outgroup (*R*) are described on the right box.

To relate our findings to different biological data, Green et al. 2010 used the D-statistic to detect introgression between Neanderthal and modern humans and had a *R* value equal to 0.873: 825,000 years separated modern humans from Neanderthal, while 6.5 million years separated humans from chimps (the outgroup) (Green et al. 2010). The study of bears of Barlow et al. (2018) used two different outgroups, black bears and pandas, with *R* values *ca*. 0.4 and 0.9, respectively. According to our simulations, all these values fall within the range of high probability of erroneous interpretation.

#### 3.2.3 The distance between donor and recipient

Species that are genetically close have a higher chance to introgress, which could mitigate the previous result. Indeed, if the distance between outgroup and ingroup is sufficient to prevent introgression between the two, then putative midgroups ghost lineages may also be too distant. It is well known that the probability of hybridization, and consequently of introgression, decreases as genetic distance between species increases (Edmands 2002; Mallet 2005; Chapman and Burke 2007; Montanari et al. 2014).

To test whether this observation mitigates the importance of ghost lineages when detecting introgression, we used different values of *α*, a parameter that lets the probability of introgression vary with the phylogenetic distance between donor and recipient (see Material and Methods). We used *α* = {0, 1, 10, 100, 1000}. When *α* = 0, introgressions occur uniformly at random; when *α* = 1000, introgressions occur almost exclusively between sister taxa (Figure 6A).

**Figure 6.**
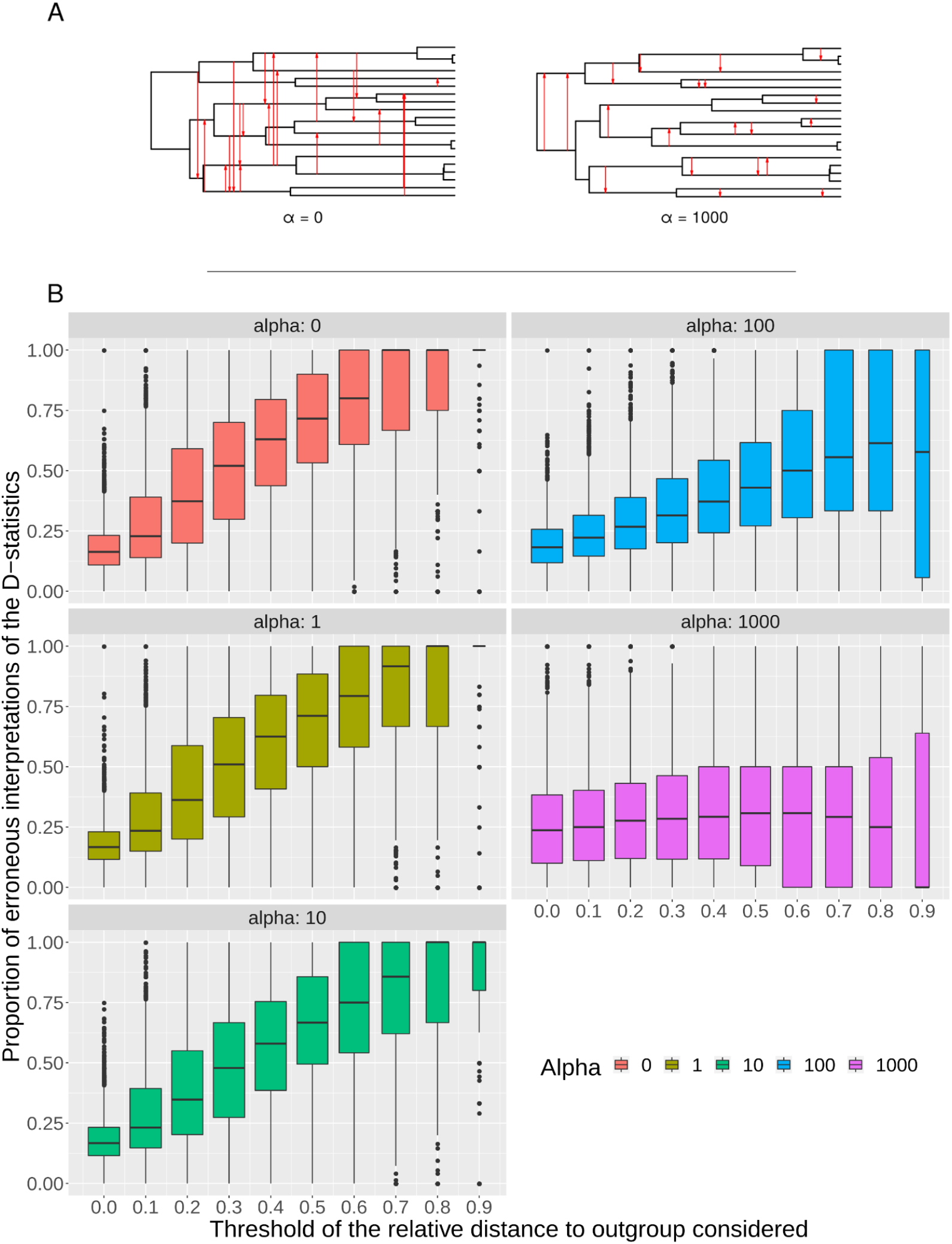
The effect of the probability of introgression on the proportion of erroneous interpretations of the D-statistic. **A.** Illustration of the effect of the *α* parameter, which imposes constraints on introgression in relation to phylogenetic distance. **B**. Relationship between relative outgroup distance (*x*-axis) and the proportion of erroneous interpretations (*y*-axis) for different levels of constraint on introgression related to phylogenetic distance, determined by *α* = {0, 1, 10, 100, 1000}.

We observed that the impact of outgroup distance on the proportion of erroneous interpretations decreased with increasing values of *α* (Figure 6B). As expected, in simulations where alpha is maximum (*α* = 1000), the proportion of significant D-statistics due to ghost introgressions is not affected by the distance separating ingroup and outgroup. Nevertheless, this proportion remains quite high, and its median value does not fall below 25% under our settings. For other values of *α*, this proportion is higher and increases with distance to the outgroup.

We investigated what could be a realistic value of alpha in biological data. We used phylogenetic trees from several studies where one or several introgression paths were identified. We counted the number of internal nodes between donor and recipient. Then, we randomly simulated the same number of introgressions with different values of alpha and calculated the average number of internal nodes between donor and recipient (removing introgressions between sister branches as this cannot be observed when using D-statistic). We retained the value of alpha giving the average number of internal nodes that was closest to what is observed in the biological data (Supplementary Material Section 4). We analyzed the bear phylogeny of Hailer et al. (2012) and found that *α* = 0 gives the closest result (actual number of nodes = 5; with *α* = 0, the average number of nodes from the simulations was 4.5). In the phylogeny of the *Bos* species complex of Wu et al. (2018), the actual number of nodes is higher than in our simulations even with *α* = 0. The same result was found with the phylogeny of the *Anopheles gambiae* species complex of Fontaine et al. (2015). By contrast, for the woodcreeper phylogeny of Pulido-Santacruz et al. (2020), we found that the value closest to the biological dataset was obtained with *α* = 100. Lastly, it was estimated that values of *α* between 100 and 1000 best fit the spider phylogeny of Leduc-Robert and Maddison (2018). These results are described in full in Supplementary Materials section 3. These examples, taken from very diverse organisms, tend to show that the higher probability of introgression between closely related species is not sufficient to secure a safe zone for the ABBA-BABA test.

## 4 Discussion

### 4.1 Ghost lineages: An important factor affecting introgression tests

Different parameters are known to affect the robustness or sensitivity of the D-statistic. For instance, variations in population size (Eriksson and Manica 2012; Lohse and Frantz 2014; Martin et al. 2015; Zheng and Janke 2018) and/or ancestral population structure (Durand et al. 2011; Lohse and Frantz 2014; Martin et al. 2015) have been shown to produce significant D-statistics in the absence of introgression. Applying the D-statistic to smaller genomic windows, rather than over the entire genome, gives very variable estimates of D (Martin et al. 2015). Complex introgression scenarios, with more than one introgression in the quartet, are another source of error (Rogers and Bohlender 2015; Elworth et al. 2018). Our findings suggest that, in addition to the variables listed above, the interpretation of the D-statistic should systematically, and maybe primarily, take into account ghost lineages.

Recently, Hibbins and Hahn (2021) published results that are in line with our findings. Their simulation study confirms that introgression from a midgroup ghost lineage can result in a significant D-statistic, which may lead to the misidentification of the identity of the lineages involved in the introgression if only known lineages are considered. Similar results have been observed with the D3 statistic (Hahn and Hibbins 2019), a test for detecting introgression that uses only three lineages and the branch lengths of the phylogeny. While the study of Hibbins and Hahn (2021) confirms that ghost introgressions may lead to erroneous interpretations, they do not quantify the extent to which this factor affects the interpretation of the D-statistic.

### 4.2. An intractable incompleteness

Although some families are believed to be extensively described (Chapman and Burke 2007), it is not possible and probably will never be possible to assume that we work with an exhaustive taxon set. A study from 2011 estimated that 8.7 million of eukaryote species are alive today (Mora et al. 2011), and a study from 2016 estimated that there are 1 trillion species on Earth (Locey and Lennon 2016). By contrast, 2.5 million species have been described and catalogued in The Catalogue of Life (CoL). The kingdom Animalia has the most descriptions with 1.4M catalogued species, while Plantae, Fungi and other kingdoms have 375K, 145K and 81K catalogued species, respectively. This means that, at best, we know 25% of the biodiversity that is alive today. In practice, there is a strong disparity in the percentage of undescribed species among taxonomic groups. Chapman estimated in 2009 that less than 20% of the phylum Chordata is yet to be described while 80% of the class Insecta is still unknown to us (Chapman et al. 2009). The proportion of undescribed species is even higher in prokaryotes with less than 1% of species described for viruses, Archaea and Bacteria, and because microbial biodiversity is harder to estimate than that of macro-organisms, these numbers could be orders of magnitude lower. Thus, the effects shown here cannot be circumvented by adding or expecting more species and improving computational techniques to handle larger datasets. We are bound to work with a very small fraction of what exists. According to our simulations, with 25% of species sampled, on average more than 50% of introgressions could be due to ghost lineages and subsequently be misinterpreted by a D-statistic. This implies that, with the exception of some well described eukaryote groups such as the genus *Homo,* our lack of taxonomic knowledge will greatly impact the reliability of the D-statistic.

### 4.3 Other introgression detection methods

Several other methods have been developed to mitigate some of the limitations of the D-statistic, but their robustness to ghost lineages has not yet been explored.

It is possible to apply the D-statistic test in datasets with more than four species by performing multiple tests on different quartets. The D-statistics are then analyzed together, using the interpretation of each individual test as a constraint for the interpretation of the other tests. This enables a finer detection of introgression, the identification of donors and recipients (while a single test cannot distinguish the donor from the recipient), and possibly assigning introgression events to groups of taxa instead of single taxa (Pease et al. 2016; Rouard et al. 2018; Suvorov et al. 2020). However, if each individual test is interpreted, as it is usually done, without considering the possibility of ghost introgressions, the joint interpretation of multiple tests will miss a high number of scenarios. Moreover, there is no method that formalizes the constraints from multiple tests and no guarantee that the result is correct or unique, and that the order in which single tests are analysed does not matter.

Extensions of the D-statistic, namely the partitioned-D (Eaton and Ree 2013) and D_FOIL_ (Pease and Hahn 2015) tests, have been proposed to infer introgression in 5-taxon phylogenies (instead of four) and to polarize (in some cases a direction to the introgression can be assessed) the introgression. Although D_FOIL_ can detect more introgression patterns than ABBA-BABA, it is still blind to ghost lineages, and thus presents similar theoretical possibilities of misinterpretation as the ABBA-BABA test (see Supplementary Material Section 5 for a listing of the patterns that lead to misinterpretation). This possibility is mentioned by Pease and Hahn (2015) but the proportion of misinterpretations remains to be quantified, and alternative interpretations handling ghost lineages has not been written.

Soraggi et al. (2018) proposed an extension of the D-statistic (D_*ext*_) to study introgression events among non-African human populations using Africans as the outgroup. This test is robust to introgression from an external group which is not part of the analysed populations. However, the use of this version of D-statistic is restricted to extinct clades for which introgression events with extant species have been already identified as the Neanderthal introgression. This precludes its use in cases where there is no *a priori* knowledge on the existence of ghost lineages.

Other more global methods, such as STRUCTURE/ADMIXTURE (Pritchard et al. 2000; Tang et al. 2005), Treemix (Pickrell and Pritchard 2012) and Phylonet (Than et al. 2008; Wen et al. 2018), have been designed to detect introgression across an entire phylogeny. These tools do not consider ghost lineages and their potential effect on the detected signal. Thus, introgression events are only inferred between known lineages and branches in a tree. It is interesting to note that two of these tools, STRUCTURE and ADMIXTURE, which are popular choices for reconstructing genetic history and testing admixture scenarios, have recently been shown to be subject to misinterpretation due to ghost introgressions (Lawson et al. 2018). The impact of ghost lineages on the detection of introgression is therefore not just a question of using the right tool.

### 4.4. Application of the D-statistic in the light of ghost introgressions

Now that the importance of the possibility of ghost introgression is recognized, and that no current method is able to handle the effects of ghost lineages, it is time to adapt all methods or develop new methods to take this factor into account.

For the single D-statistic test (four-taxon quartet), the solution is simply to take this uncertainty into account by considering alternative scenarios with at least equal probability. In phylogenies with more than four taxa, the D-statistics from all quartets could be analyzed using an algorithmic method that would combine a set of scenarios. This will require formalizing the objective (*i.e*. minimizing the number of incoherences between quartet results), choosing a set of quartets according to this objective and devising a combinatorial algorithm that handles, for each quartet, information from several possible scenarios including ghost lineages.

Note that this approach will not only avoid interpretation errors, but will possibly point to the existence of unknown lineages that have contributed through introgression to the genomes of known lineages. Therefore, this approach would combine the detection of introgression and the detection of unsampled or extinct taxa. This has already been achieved with *ad hoc* methods for human (Prüfer et al. 2014; Dannemann and Racimo 2018) and whale lineages (Foote et al. 2019) and could be generalized to enrich the phylogeny of known species with unknown species, for which we have no trace other than the genes that have lived, for a while, in them. This is a promising route for future work.

## 5. Conclusion

The D-statistic is a key tool for studying introgression as it provides, in specific cases, a robust test for detecting gene flow. However, our results show that one important caveat of this test is its lack of consideration of ghost lineages, which can lead to the misinterpretation of a significant result. Thus, the *bona fide* interpretation of a single significant D-statistic should be a set of possible scenarios that include the possibility of ghost introgressions, which are equally likely in the absence of other information. Based on our simulations, we have suggested that ghost introgressions are often the most likely scenario. It is possible that in the future, the usual interpretation of a significant D-statistic, *i.e*. ingroup introgression, becomes the exception rather than the rule.

## Acknowledgments

This work was supported by the French National Research Agency (Grants ANR-18-CE02-0007-01 and ANR-19-CE45-0010). Simulations were performed using the computing facilities of the CC LBBE/PRABI. We thank Gergely Szöllosi for useful discussions

## Software Availability

All codes used to generate and analyze the simulations performed in this study are available at: https://github.com/theotricou/Ghost_abba_baba.

