## Supplementary Material for "Ghost lineages highly influence the interpretation of introgression tests"

### 1. *Equivalence between D-statistics on sequence simulations and gene tree species tree comparisons*

The standard simulation procedure to evaluate gene flow inference methods is to draw a species tree, choose an introgression between two of its branches and then simulate sequences along this reticulated species tree. ABBA-BABA patterns are read on these sequences and the tests are realized from these patterns.

We explore here a way to bypass the sequence simulation, which is the most computationally demanding. We construct a *gene tree* from the species tree and the introgression by performing a subtree prune and regraft (SPR). Then for each quartet of species (((P1,P2),P3),P4) we check if the gene tree has this topology on this quartet or an alternative one. If the gene tree has (((P1,P3),P2),P4), we identify this situation with an excess of BABA patterns when simulating sequences, and if the gene tree has (((P2,P3),P1),P4), with an excess of ABBA patterns. We discard the cases of introgressions involving P4.

We hypothesize that each time the gene tree shows one of these patterns, we would have obtained a significant D-statistic test if we had simulated sequences, with a stochastic uncertainty due to the random process and the simulation parameters, such as population size and sequence size. In order to test this statement we simulated sequences on a control dataset. For this we used *ms* (Hudson 2002) to simulate $10^6$ independent loci with a single mutation each (resulting in $10^6$ SNP pattern), evolving in populations of fixed size (Ne) of 100,000 individuals over $10^6$ generations. For each quartet with topology (((P1,P2),P3),O) in the species tree and for each introgression simulated, we counted the number of loci with ABBA and BABA patterns. We then computed the D-statistics. The R package *Coala* (Staab and Metzler 2016) was used to execute *ms* and to analyse the resulting data.

In Supplementary Figure 1 we show that the proportion of erroneous interpretation is highly correlated for both approaches ($R^2 > 0.96$).

In consequence we kept only the computationally less intensive approach for the study with several parameters to explore.

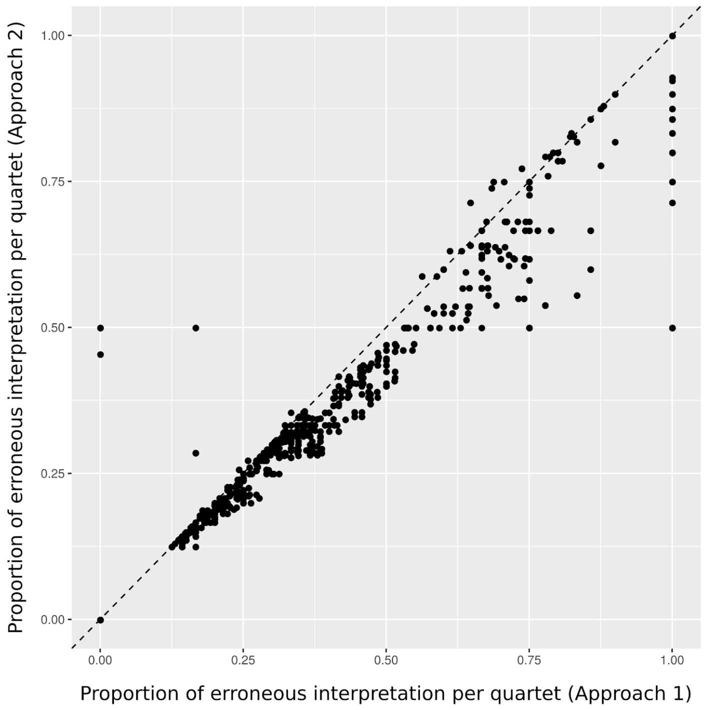


**Supplementary Figure 1.** Comparison of the simulation procedure we used and a more complete and more computationally demanding one: computation of the D-statistics based on simulated SNPs (x-axis) or inference based on the topology of the trees after introgression (y-axis). The proportion of erroneous interpretation over 500 introgressions for a unique species tree (*N*=40, *α*=0, extinction probability *pex*=0.5) was computed for each quartet (black dots). The dashed line represents the first diagonal. The squared Fisher correlation coefficient is $R^2 = 0.9618662$, providing a robust basis for using one approach as a proxy for the other

### 2. *All results of varying parameters*

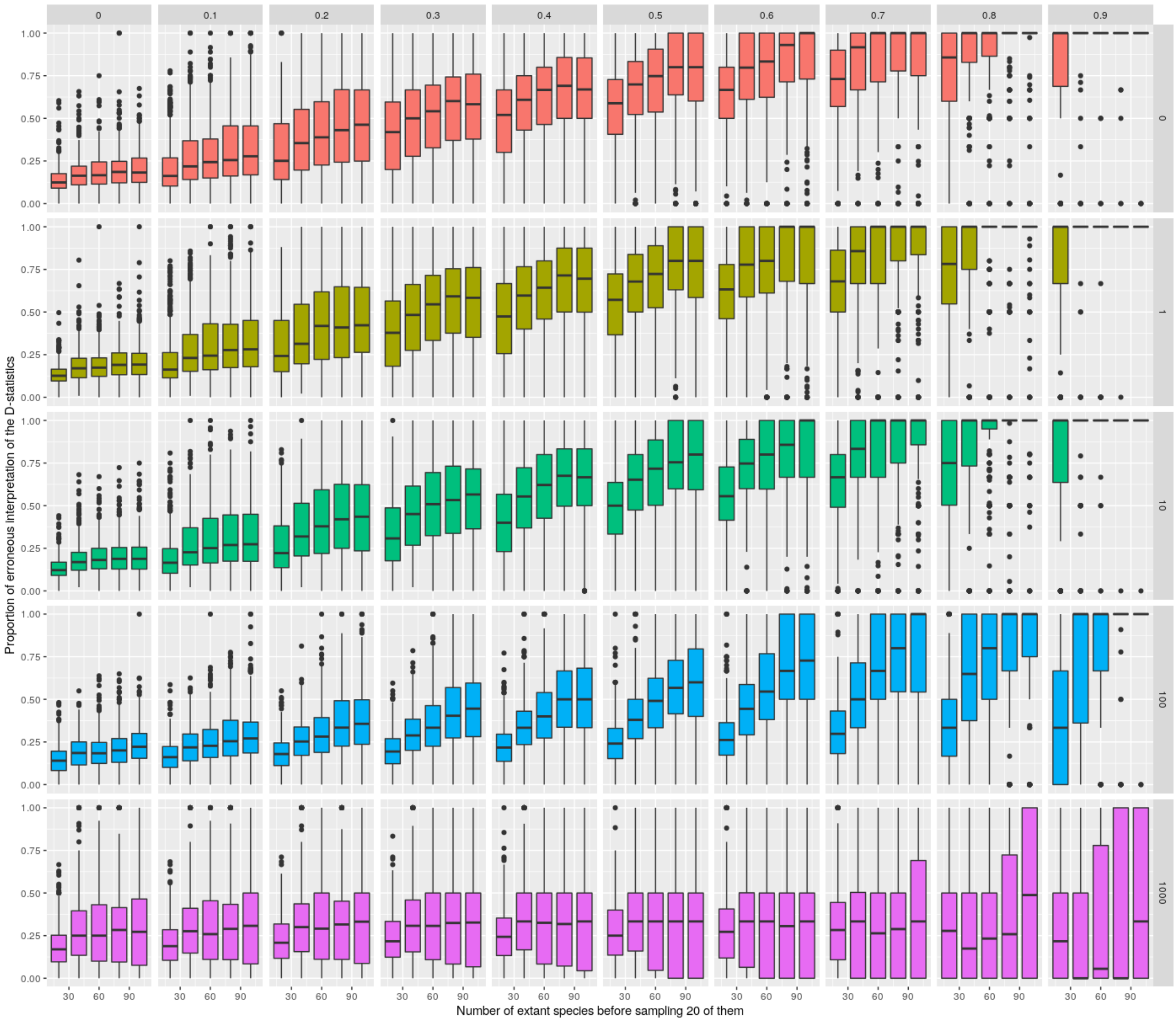


**Supplementary Figure 2.** Proportion of erroneous interpretation of D-statistics for all subsets of parameters tested. Mean proportion of erroneous interpretations observed (**y-axis**) as a function of the taxonomic sampling effort (x-axis) 20 species sampled for N extent species simulated (N = 20, 40, 60, 80, 100) by the distance to the outgroup (**columns**) for different strengths of the phylogenetic distance effect (**lines**) as controlled by α (α = 0, 1, 10, 100 and 1000).

#### *3. No effect of the number of extinct species*

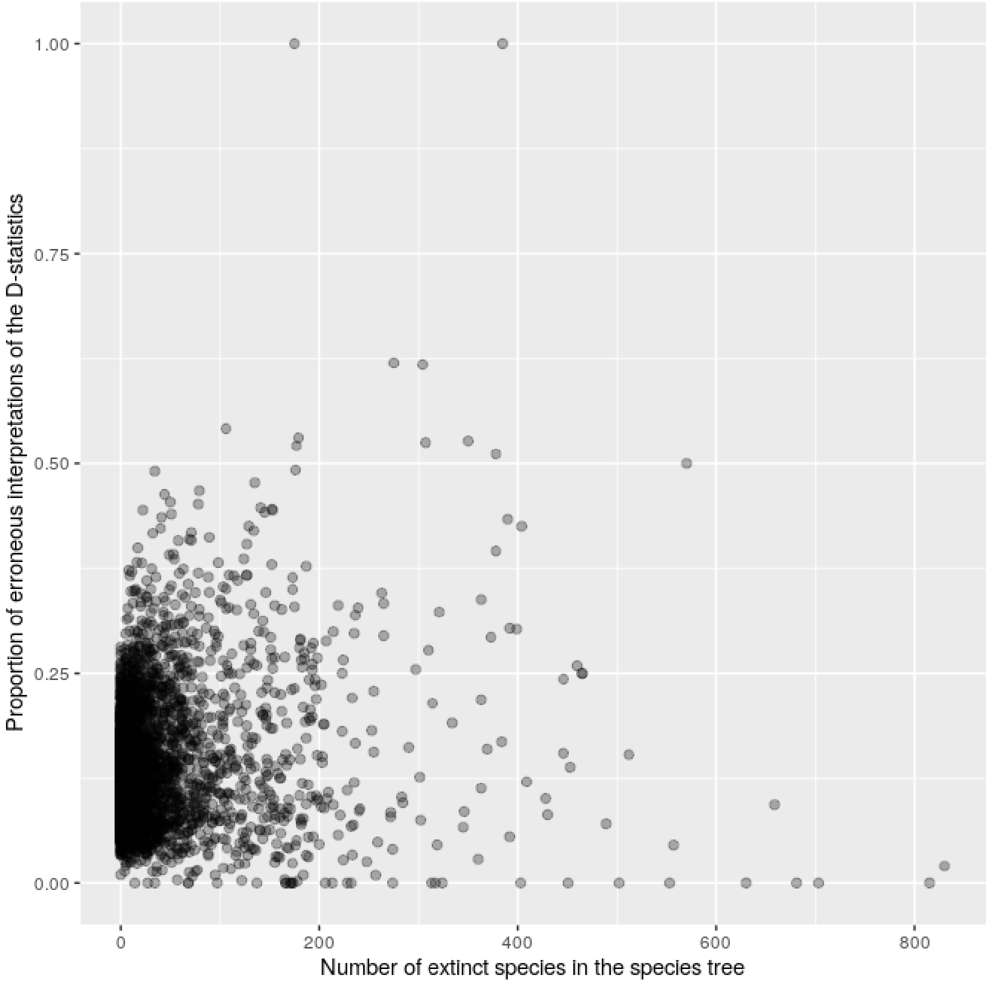


**Supplementary Figure 2.** Effect of the number of extinct species on the proportion of erroneous interpretation of the D-statistics. Proportion of erroneous interpretations observed (y-axis), function of the number of extinct lineages in the species tree (x-axis). Species trees were simulated using 4 extinction rates, pext (pext = 0, 0.3, 0.6, 0.9). For each parameter pext, 1000 species trees were simulated with 20 extant species at the end and with 100 introgressions sampled.

### 4. *Estimating alpha on biological data*

In the five following figures, five different phylogenies were taken from the literature :

(A) the bear phylogeny from Hailer et al. (2012), Figure 1A in this paper

(B) the bos phylogeny from Wu et al. (2018), Figure 1 and Supplementary Figure 13 in this paper

(C) the mosquito phylogeny from Fontaine et al. (2015), Figure 1C in this paper

(D) the woodcreeper phylogeny from Pulido-Santacruz, Aleixo, and Weir (2020), Figure 5 in this paper

(E) the spider phylogeny from Leduc-Robert and Maddison (2018) Figure 1 in this paper

For each phylogeny, there are a certain number of introgressions documented. We simulated the same number of introgressions with alpha = {0,1,10,100,1000}. For the observed and simulated situations we computed the mean number of nodes in the phylogeny separating the donors and recipients of introgressions, excluding introgressions between sister branches in the simulations (because they cannot be detected in the biological data). This mean number is drawn on the five figures, in function of alpha, and the observed number is added on the figure by an horizontal line.

This shows ranges of alpha parameters that can best explain the observations. This range highly depends on the dataset, and shows that there is no unique possible choice of alpha. In consequence the range in which we make this parameter vary is not unrealistic, and no extremal value can be a priori prefered.

**Supplementary Figures 3-7**

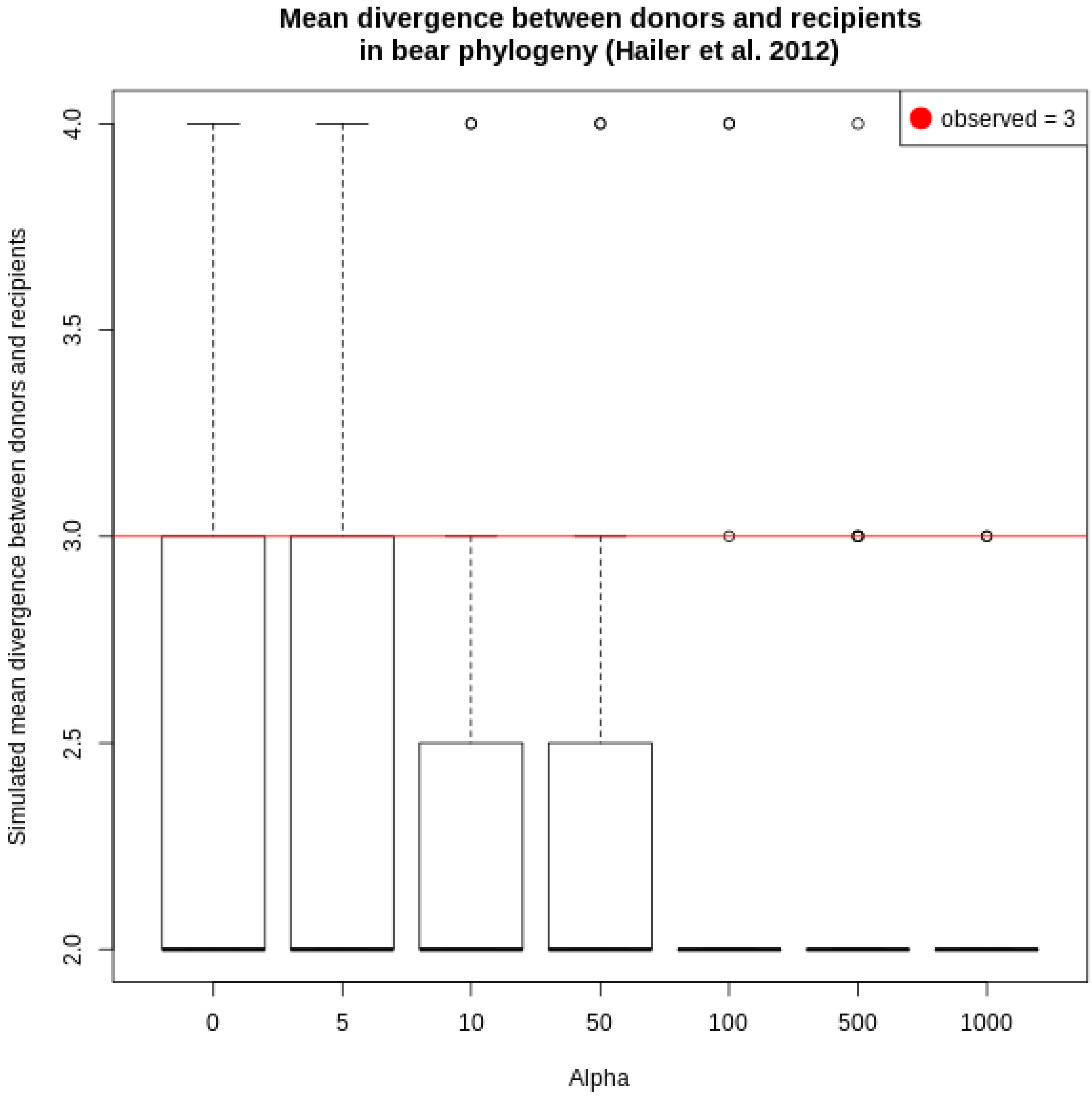

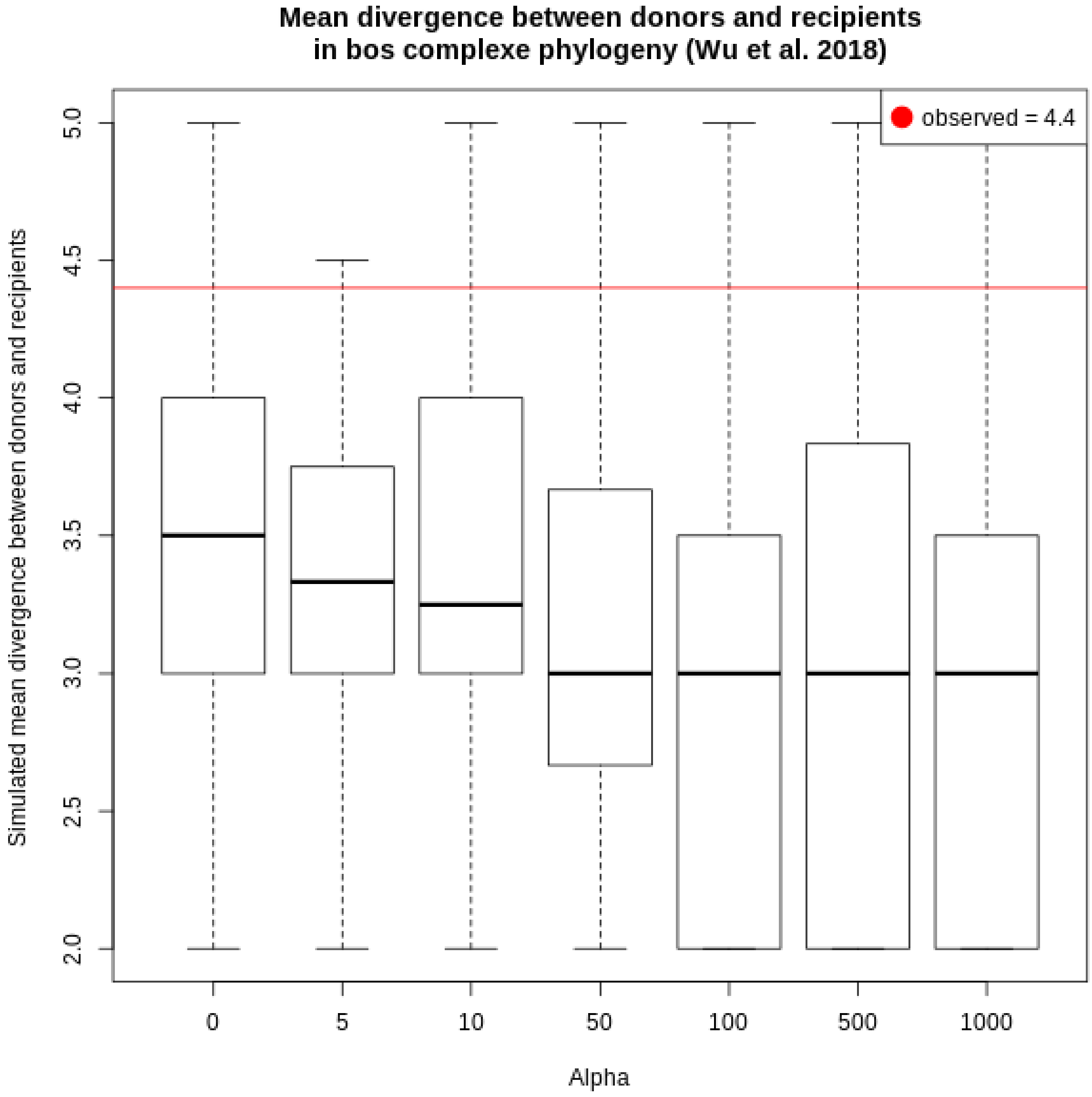

Mean divergence between donors and recipients
in bos complexe phylogeny (Wu et al. 2018)
observed = 4.4
Simulated mean divergence between donors and recipients
5.0
4.5
4.0
3.5
3.0
2.5
2.0
0
5
10
50
100
500
1000
Alpha

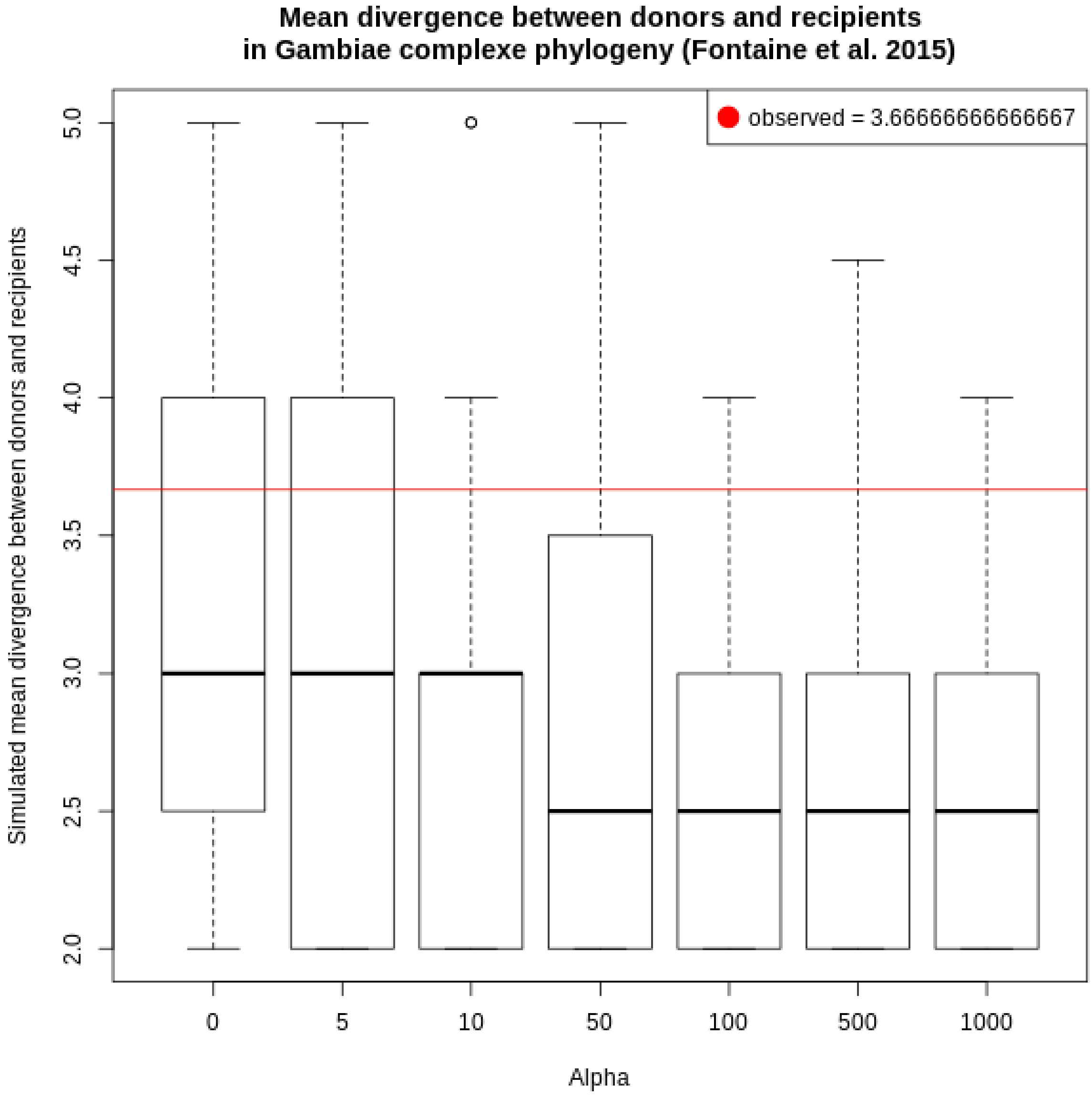

Mean divergence between donors and recipients
in Gambiae complexe phylogeny (Fontaine et al. 2015)
observed = 3.66666666666667
Simulated mean divergence between donors and recipients
2.0
2.5
3.0
3.5
4.0
4.5
5.0
0
5
10
50
100
500
1000
Alpha

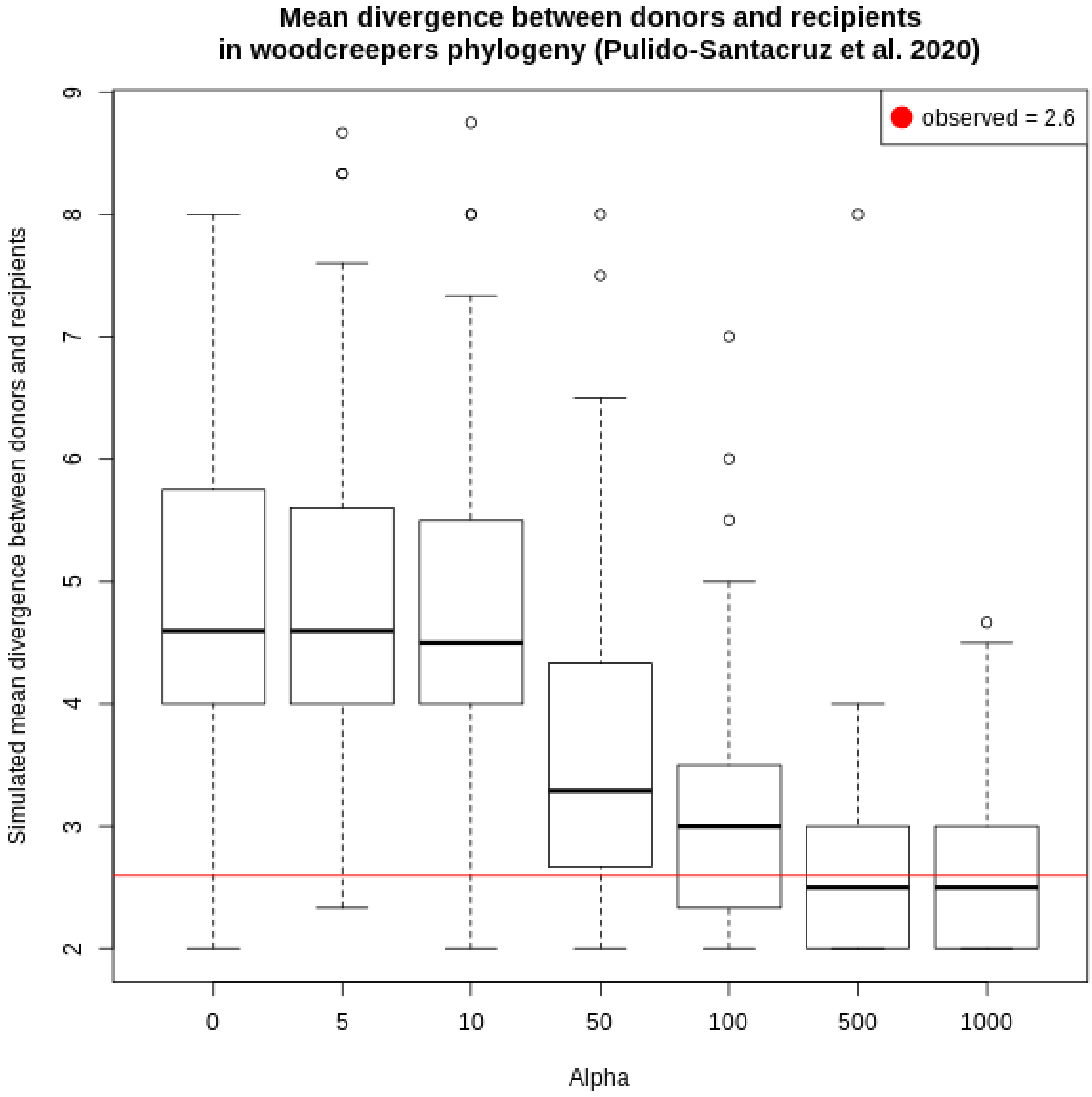

Mean divergence between donors and recipients
in woodcreepers phylogeny (Pulido-Santacruz et al. 2020)
observed = 2.6
Simulated mean divergence between donors and recipients
2
3
4
5
6
7
8
9
0
5
10
50
100
500
1000
Alpha

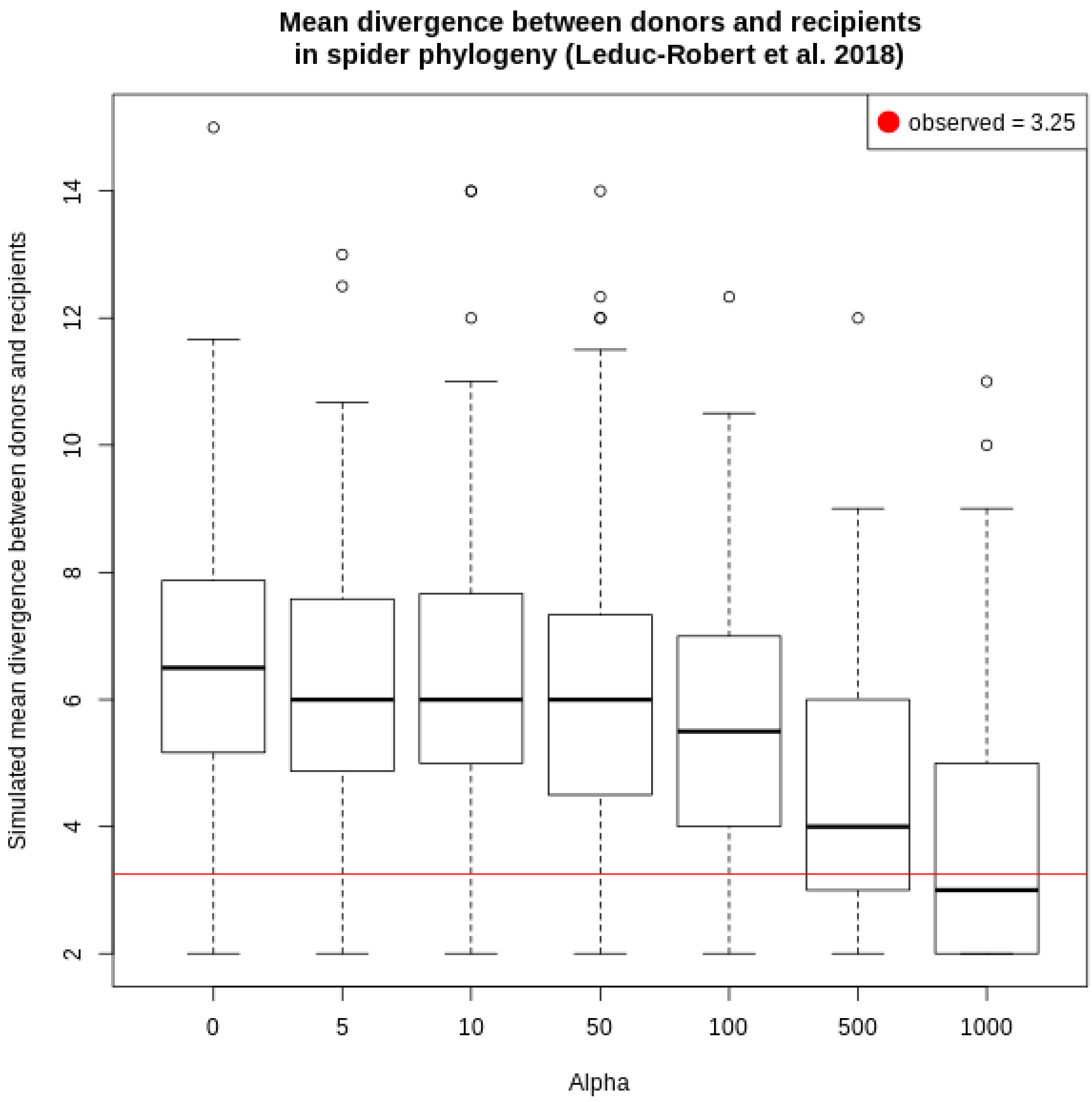
Mean divergence between donors and recipients
in spider phylogeny (Leduc-Robert et al. 2018)
observed = 3.25
Simulated mean divergence between donors and recipients
2
4
6
8
10
12
14
0
5
10
50
100
500
1000
Alpha

### 5. *Erroneous interpretations in the $D_{FOIL}$ test*

In the main text we argue that just as the D-statistic, $D_{FOIL}$ is subject to misinterpretations. We detail this statement here, and need to recall first how it is computed.

*2.1. How the $D_{FOIL}$ is computed*

For a combination of 5 lineages with symetrical topology (((P1,P2),(P3,P4)),O), the 4 D-like statistics with the following equations are computed (Pease and Hahn 2015):

$$D_{FO}= \frac{(\Sigma BABAA + \Sigma BBBAA + \Sigma ABABA + \Sigma AAABA) - (\Sigma BAABA + \Sigma BBBAA + \Sigma ABBAA + \Sigma AABAA)}{(\Sigma BABAA + \Sigma BBBAA + \Sigma ABABA + \Sigma AAABA) + (\Sigma BAABA + \Sigma BBBAA + \Sigma ABBAA + \Sigma AABAA)}$$

$$D_{IL}= \frac{(\Sigma ABBAA + \Sigma BBAAA + \Sigma BAABA + \Sigma AAABA) - (\Sigma ABABA + \Sigma BBABA + \Sigma BABAA + \Sigma AABAA)}{(\Sigma ABBAA + \Sigma BBAAA + \Sigma BAABA + \Sigma AAABA) + (\Sigma ABABA + \Sigma BBABA + \Sigma BABAA + \Sigma AABAA)}$$

$$D_{FI}= \frac{(\Sigma BABAA + \Sigma BABBA + \Sigma ABABA + \Sigma ABAAA) - (\Sigma ABBAA + \Sigma ABBBA + \Sigma BAABA + \Sigma BAAAA)}{(\Sigma BABAA + \Sigma BABBA + \Sigma ABABA + \Sigma ABAAA) + (\Sigma ABBAA + \Sigma ABBBA + \Sigma BAABA + \Sigma BAAAA)}$$

$$D_{OL} = \frac{(\Sigma BAABA + \Sigma BABBA + \Sigma ABBAA + \Sigma ABAAA) - (\Sigma ABABA + \Sigma ABBBA + \Sigma BABAA + \Sigma BAAAA)}{(\Sigma BAABA + \Sigma BABBA + \Sigma ABBAA + \Sigma ABAAA) + (\Sigma ABABA + \Sigma ABBBA + \Sigma BABAA + \Sigma BAAAA)}$$

Binomial tests are performed to evaluate whether the difference between both elements framing the minus sign of each equation was significant, in order to assign a "+", "-" or "0" sign. In (Pease and Hahn 2015), 8 unique patterns of $D_{FOIL}$ were linked to different polarized introgression events (with an explicit direction) and another 2 to pairs of non polarized (both directions) events (see Table 1. in Pease and Hahn 2015).

*2.2. $D_{FOIL}$, a 5-taxon extension of D-statistics, rarely solves the issue raised by ghost introgressions.*

If we examine the DFOIL statistic with the possibility of presence of ghost lineages, we can observe two additional $D_{FOIL}$ patterns, "00++" and "00--" that can be interpreted as non polarized events. Furthermore, any non polarized event can be explained either by an introgression between an ancestor lineage of one clade and a species from the opposite clade or an introgression from a midgroup ghost lineage to the second species of the opposite clade. For example, the Dfoil pattern "++00" arises from the event P1P2<->P3 but could also be observed following the event Ghost->P4. For "--00", events are P1P2<->P4 and Ghost->P3. For the two new patterns, "00--" and "00++", events are P3P4<->P1 and Ghost->P2 and events are P3P4<->P2 and Ghost->P1 respectively. It should be noted that, similarly to the D-statistic, an introgression from the outgroup lineages or an external lineages to the quintet will produce the same pattern as a midgroup ghost interpretation. Given that the ancestor of P3 and P4 is always older than the ancestor of P1 and P2, this implies that a lineage with no descendant available inside the ingroup is the donor for those two $D_{FOIL}$ patterns, either a sister lineage to P3P4 or a midgroup ghost lineage. Conversely polarized events can not be explained by any events involving midgroup ghost lineages. This means that Dfoil can only be erroneously interpreted if the pattern is non polarized.


Fontaine, Michael C., James B. Pease, Aaron Steele, Robert M. Waterhouse, Daniel E. Neafsey, Igor V. Sharakhov, Xiaofang Jiang, et al. 2015. "Extensive Introgression in a Malaria Vector Species Complex Revealed by Phylogenomics." *Science* 347 (6217): 1258524. https://doi.org/10.1126/science.1258524.

Hailer, Frank, Verena E. Kutschera, Björn M. Hallström, Denise Klassert, Steven R. Fain, Jennifer A. Leonard, Ulfur Arnason, and Axel Janke. 2012. "Nuclear Genomic Sequences Reveal That Polar Bears Are an Old and Distinct Bear Lineage." *Science* 336 (6079): 344–47. https://doi.org/10.1126/science.1216424.

Hudson, R. R. 2002. "Generating Samples under a Wright-Fisher Neutral Model of Genetic Variation." *Bioinformatics* 18 (2): 337–38. https://doi.org/10.1093/bioinformatics/18.2.337.

Leduc-Robert, Geneviève, and Wayne P. Maddison. 2018. "Phylogeny with Introgression in Habronattus Jumping Spiders (Araneae: Salticidae)." *BMC Evolutionary Biology* 18 (1): 24. https://doi.org/10.1186/s12862-018-1137-x.

Pease, James B., and Matthew W. Hahn. 2015. "Detection and Polarization of Introgression in a Five-Taxon Phylogeny." *Systematic Biology* 64 (4): 651–62. https://doi.org/10.1093/sysbio/syv023.

Pulido-Santacruz, Paola, Alexandre Aleixo, and Jason T. Weir. 2020. "Genomic Data Reveal a Protracted Window of Introgression during the Diversification of a Neotropical Woodcreeper Radiation*." *Evolution* 74 (5): 842–58. https://doi.org/10.1111/evo.13902.

Staab, Paul R., and Dirk Metzler. 2016. "Coala: An R Framework for Coalescent Simulation." *Bioinformatics* 32 (12): 1903–4. https://doi.org/10.1093/bioinformatics/btw098.

Wu, Dong-Dong, Xiang-Dong Ding, Sheng Wang, Jan M. Wójcik, Yi Zhang, Małgorzata Tokarska, Yan Li, et al. 2018. "Pervasive Introgression Facilitated Domestication and Adaptation in the Bos Species Complex." *Nature Ecology & Evolution* 2 (7): 1139–45. https://doi.org/10.1038/s41559-018-0562-y.
